# Alkane and wax ester production from lignin derived molecules

**DOI:** 10.1101/502468

**Authors:** Milla Salmela, Tapio Lehtinen, Elena Efimova, Suvi Santala, Ville Santala

## Abstract

Lignin has potential as a sustainable feedstock for microbial production of industrially relevant molecules. However, the required lignin depolymerization yields a heterogenic mixture of aromatic monomers that are challenging substrates for the microorganisms commonly used in industry. Here, we investigated the properties of lignin-derived molecules (LDMs), namely coumarate, ferulate, and caffeate, in the synthesis of biomass and products in a LDM-utilizing bacterial host *Acinetobacter baylyi* ADP1. The biosynthesis products, wax esters and alkanes, are relevant compounds for the chemical and fuel industries. In *A. baylyi* ADP1, wax esters are produced by a native pathway, whereas alkanes are produced by a synthetic pathway introduced to the host. Using individual LDMs as substrates, the growth, product formation, and toxicity to cells were monitored with internal biosensors. Of the tested LDMs, coumarate was the most propitious in terms of product synthesis. Wax esters were produced from coumarate with a yield and titer of 40 mg /g_coumarate_ and 221 mg/L, whereas alkanes were produced with a yield of 62.3 μg /g_coumarate_ and titer of 152 μg/L. This study demonstrates the microbial preference for certain LDMs, and highlights the potential of *A. baylyi* ADP1 as a convenient host for LDM upgrading to value-added products.

## Introduction

Microbial processes utilizing non-edible biomass as a substrate can offer a sustainable solution for the production of fuels and chemicals. Comprehensive utilization of cheap waste streams obtained from agriculture and forest industry, could improve the economic viability of the bioprocesses (FitzPatrick et al., 2010), (Elshahed, 2010), (Clark et al., 2006), (Steen et al., 2010)(Peralta-Yahya et al., 2012). Currently lignin, one of the most abundant biopolymers, is underutilized mainly due to its recalcitrance and heterogeneity. Because of the complex structure, feedstock originating from lignocellulose and lignin treatment processes contain a heterogeneous mixture of compounds, including phenols, acids and residual sugars (Katahira et al., 2016)(Constant et al., 2016)(Abdelkafi et al., 2011)(Sun et al., 2015)(Li et al., 2012) (Karp et al., 2016) (Raj et al., 2007). Many lignin-derived molecules (LDMs) are growth inhibitors poorly tolerated by the most commonly used microbial production hosts, such as *Escherichia coli* and *Saccharomyces cerevisiae*. (Rumbold et al., 2009)(Sun et al., 2001)(Palmqvist and Hahn-Hägerdal, 2000)(Jönsson and Martín, 2016)(Adeboye et al., 2014). More importantly, these strains lack the catabolic pathways for LDM utilization. Thus, increasing attention is given towards microbial hosts that are capable of tolerating and metabolizing the LDMs, and can be employed as modular cell factories for the synthesis of products of interest (Nielsen et al., 2009)(Freed et al., 2018).

*Acinetobacter baylyi* ADP1 is an example of a wide substrate range bacterium that can degrade and utilize LDMs for growth and biosynthetic pathways. In *A. baylyi* ADP1, aromatic compounds are channeled to the central metabolism *via* a β-ketoadiapate pathway (Bleichrodt et al., 2010). In this upper funneling pathway, structurally different compounds are converted to central intermediates, protocatechuate or catechol, before entering the β-ketoadiapate pathway and eventually resulting in common metabolites acetyl-CoA and succinyl-CoA (Ornston, 1966)(Fuchs et al., 2011) (Fischer et al., 2008). Thus, the β-ketoadiapate pathway could be exploited in the synthesis of a broad range of acetyl-CoA derived compounds from lignin-derived feedstock. Previous demonstrations of microbial upgrading of LDMs include the production of medium-chain (C_6_-C_14_) polyhydroxyalkanoates (PHA) by *Pseudomonas putida* (Linger et al., 2014) and triacylglycerols by *Rhodococcus opacus* (Kosa and Ragauskas, 2012). Lignin-derived phenolic compounds such as coumarate, ferulate and caffeate are structural analogues metabolized through the protocatechuate branch of the upper funnelling pathway (Fischer et al., 2008). The occurrence and position of substitution groups in the aromatic ring may affect the biochemical reactions and inhibitory effects of these compounds. On the other hand, the diversity of phenolic compounds released from biomass depends on the chosen pre-treatment method and the origin of the biomass (Constant et al., 2016). Thus, studies on the substrate preferences of the microbial cell factories promotes preferable choice of biomass and pretreatment methods.

In addition to LDM utilization, A. *baylyi* ADP1 has interesting features of being readily genetically engineered organism (Metzgar, 2004) (Elliott and Neidle, 2011) (de Berardinis et al., 2008) that also accumulates industrially relevant long-chain alkyl esters (wax esters). Similarly to other storage lipids, wax esters are produced intracellularly in nitrogen-deficient conditions with excess carbon reserves (Alvarez and Steinbüchel, 2003)(Fixter et al., 1986)(Santala et al., 2014). The wax esters produced by *A. baylyi* ADP1 resemble the structure of jojoba-oil (produced by *Simmondsia chinensis)* with a typical carbon content of C_32_-C_36_ (Fixter et al., 1986)(Kalscheuer and Steinbüchel 2003) (Lehtinen et al., 2018a), The composition of wax esters can be modified by alternating process conditions (Dewitt et al., 1982) or by genetically rewiring pathways (Santala et al. 2014), presenting further opportunities in product tailoring. On the other hand, genome integrated synthetic pathways provide practical means to produce non-native products of industrial relevance. For example, we have previously constructed an *A. baylyi* ADP1 strain that accumulates intracellular alkanes by expressing a non-native fatty acid reductase (AAR) and an aldehyde deformylating oxygenase (ADO) (Lehtinen et al., 2017b). In the strain, the naturally occurring alkane degradation and wax ester synthesis pathways of *A. baylyi* ADP1 were disrupted by targeted gene knock-outs and an optimized alkane production pathway was integrated (Lehtinen et al., 2017b). Microbial wax esters and alkanes have previously been produced using carbon sources such as glucose and organic acids (Kannisto et al., 2014) (Lehtinen et al., 2018a)(Salmela et al., 2018a) (Santala et al., 2011) (Schirmer et al., 2010)(Lehtinen et al., 2017b)(Cao et al., 2016) (Fatma et al., 2018)(Lehtinen et al., 2017a).

Here, we demonstrate the production of wax esters (C_32–34_) and alkanes (C_17_) by *A. baylyi* ADP1 from LDMs, namely ferulate, caffeate, and coumarate. We profiled the growth and tolerance against these aromatic compounds, and determined how efficiently the compounds are directed to the synthesis pathways of interest. Thereafter, we utilized the native pathway of *A. baylyi* ADP1 to produce long chain alkyl esters (wax esters) from the most optimal compound (coumarate). To demonstrate the metabolic flexibility of *A. baylyi* and the potential of LDMs as a feedstock for a range of industrially relevant products, we also produced alkanes from coumarate by an engineered strain.

## Materials and methods

### Strains, media and components

*A. baylyi* ADP1 ‘sensor-strain’ (Santala et al. 2011) –designated here as the wax ester producing WP strain — was used for internal aldehyde monitoring and wax ester production. The WP strain originates from the wild type strain *A. baylyi* ADP1 (DSM 24193) with a bacterial luciferase gene iluxAB replacing gene poxB (ACIAD3381) associated with pyruvate dehydrogenase activity. The WP strain genotype is *A. baylyi* ADP1Δ*poxB*:: *luxAB*,cm^r^. A biosensor strain originating from the same wild type modified to produce alkanes instead of wax esters (Lehtinen et al. 2017a) – designated as the alkane producing AP strain — was used for internal aldehyde and alkane monitoring, and alkane production. In this strain, the gene encoding native alkane degrading activity (AlkM) of *A. baylyi* has been replaced by GFP-gene under a native alkane inducible promoter. Additionally, it has two non-native genes aar (acyl-acyl carrier protein (ACP) reductase) and ado (aldehyde-deformylating oxygenase) from *Synechococcus elongates* integrated in the genome replacing a putative prophage segment (pp2), to allow alkane synthesis. The gene expression from this synthetic alkane-producing pathway is controlled by LacI and isopropyl β-D-1-thiogalactopyranoside (IPTG) inducible promoter.

Furthermore, the strain AP has the genes ACIAD3383–3381 knocked-out and replaced by the bacterial luciferase genes *luxAB*. The ACIAD3383 (acr1) is associated with reduction of fatty acyl-CoA, thus its elimination removes the native long chain fatty aldehyde production related to the wax ester production. The genotype of AP strain is *A. baylyi*

ADP1Δ*poxB*Δ*metY*Δ*acr1*::*luxAB*,cm^r^ Δ*alkM*::*sfgfp*,kan^r^ Δpp2::*aar,ado*,spc^r^.

All cultivations were done in mineral salts media (Hartman et al. 1989) with some modifications (K_2_HPO_4_ 3.88 g/L, NaH_2_PO_4_ 1.63 g/L, (NH_4_)_2_SO_4_ 2.00 g/L, MgCl_2_.6H_2_O 0.1 g/L, EDTA 10 mg/L, ZnSO_4_.7H_2_O2 mg/L, CaCl_2_.2H_2_O 1 mg/L, FeSO_4_.7H_2_O 5 mg/L, Na_2_MoO_4_.2H_2_O0.2 mg/L, CuSO_4_.5H_2_O 0.2 mg/L, CoCl_2_.6H_2_O 0.4 mg/L, MnCl_2_.2H_2_O 1 mg/L) supplemented with 0.2% casein amino acids and antibiotics when required (kananmycin 50 μg/μL). Sodium acetate (25 mM) and coumarate, ferulate, or caffeate (15 mM) were used as carbon sources if not stated otherwise. IPTG (100 μM) (ThermoFisher, USA) was used for induction of the alkane production. All analytical compounds were purchased either form Sigma (USA) or Tokyo chemical industries (Japan).

### Comparison of different LDMs as carbon source

Growth and fatty aldehyde formation of the wax ester producing WP strain and the AP strain were measured on 96-well plates (Greiner Bio-one, Austria). The cells were incubated in a total volume of 200 μL and supplemented with 25 mM of acetate and either 15 mM of coumarate, ferulate, or caffeate. Cultivations supplemented with either 25 mM acetate or 0.2% casein amino acids were used as control cultivations. The cells were cultivated in Spark microplate reader (Tecan, Switzerland) at 30 °C for 30 hours. Luminescence and optical density (OD) at 600 nm were measured at 30 min intervals and the plate was shaken for 5 min before reading (108 rpm, 2.5 mm amplitude). Additionally, luminescence and fluorescence signals (excitation 485±10 nm, emission 510±5 nm) were measured from the AP strain. The fluorescence signal was divided by the maximum OD_600_ value to indicate produced alkanes per biomass. Media supplemented with the corresponding carbon source without inoculant were used as blanks and the background signal was subtracted from the obtained sample values. All cultivations were conducted in triplicates. The AP strain was induced with 100 μM IPTG and supplemented with kanamycin (50 μg/mL). Similar setup was used for studying substrate toxicity with the WP strain except that acetate (25–200 mM) and coumarate (13–120 mM) mixtures of various concentrations were used as carbon sources in 48 h cultivations.

### Alkane and wax ester production from coumarate in 50 ml batch cultures

The WP and AP strains were grown on 25 mM acetate and 15 mM coumarate for 24 hours in a total volume of 50 mL in 250 ml flasks at 30 °C and 300 rpm. Samples for analyses were collected after 8, 12 and 24 h of cultivation. Acetate and coumarate concentrations were measured with HPLC and cell growth (OD) was measured with spectrophotometer at 600 nm. Alkanes and wax esters were analyzed from extracted lipids with thin layer chromatography (TLC) (wax esters) or gas chromatography-mass spectrometry (GC-MS) (alkanes) as described in analytical methods. Control cultivations contained 0.2% casein amino acids or 0.2% casein amino acids and 25 mM acetate. All cultivations were done as triplicates. The C/C yield was calculated by dividing the carbon content of heptadecane with the carbon content of the substrates (acetate and coumarate) after subtracting the titer from the casein amino acid control cultivation. Similarly, the yield was calculated as g/g_coumarate_ consumed after subtracting the titer from the acetate control cultivation.

### Wax ester production from coumarate in 1-L bioreactor

Larger scale bioreactor experiment was conducted with the WP strain in a 1-liter reactor (Sartorius Biostat B plus Twin System, Germany) with an online pH and pO_2_ monitoring system and automated O_2_ feed. An initial media volume of 750 ml was supplemented with 25 mM of acetate and 15 mM coumarate and inoculated with the WP strain pre-cultivated in 100 mM acetate, the initial OD being 0.14. Temperature was set to 30 °C and stirring to 300 rpm. Samples were collected periodically either as 50 ml (for NMR, CDW, HPLC measurements) or as 5 ml (HPLC and OD measurements). Coumarate was supplemented to the reactor after carbon depletion in a total volume of 20 ml in concentrations of either 24.5 mM or 34.5 mM. The mg/g yield for wax esters were calculated with an average chain length of 34 carbons (506 g/mol) (Lehtinen et al., 2018b) from the 34.5 mM coumarate supplementation.

### Analytical methods

Acetate, coumarate and 4-HBA concentrations were measured with HPLC (Agilent 1100 series, HewlettPackard, Germany) equipped with Fast acid H+ column (Phenomex, USA), a degasser (G1322A) and an UV-detector (G1315A) using 0.005 N H_2_SO_4_ as eluent. The pump (G1211A) flow was adjusted to 1 ml/min, the column temperature to 80 °C, and peaks were identified at wavelength 310 nm for coumarate and 210 nm for acetate and 4-HBA by comparing the retention times and spectral profiles to prepared standards.

Wax esters and alkanes were extracted from cells by methanol-chloroform extraction as described previously (Santala et al., 2011): extraction was conducted for cell pellets obtained by centrifugation (12 000 g × 5 min) from equal volumes of cell cultures (12 ml). The cell pellets were suspended in methanol (500 μl), chloroform was added (250 μl), and the samples were incubated at room temperature with gentle mixing for 1 hour. Thereafter, chloroform (250 μl) and PBS (250 μl) were added, and the gentle mixing was continued for two more hours. Finally, the samples were centrifuged and the lower phase (chloroform) was used for analyses.

Thin layer chromatography (TLC) analysis was carried out with 10 × 10 cm HPTLC Silica Gel 60 F254 glass plates with 2.5 × 10 cm concentrating zone (Merck, USA). Extracted samples were loaded on the plate (30 μl) and jojoba-oil was used as a standard for wax esters. The mobile phase consisted of n-hexane:diethylether:acetic acid (90:15:1). For visualization, the plate was stained with iodine. Alkanes were measured with GC-MS (Agilent Technologies 6890N/5975B) equipped with HP-5MS 30 m × 0.25 mm column with 0.25 μm film thickness. Helium flow rate was adjusted to 4.7 ml/min and a 1 μl splitless injection was used. The oven program was adjusted to 55 °C hold 5 min, 55–280 °C 20°/min ramp and 280 °C hold 3 min. Scanning occurred at 50–500 *m/z*, 1.68 scan/s. Chromatograph peaks were identified based on the NIST library (Version 2.2/Jun 2014) and on heptadecane external standards (Sigma-Aldrich, USA). The accumulation of 8-heptadecene was determined only qualitatively by comparing the chromatograph peaks within the samples and to the GC library as a standard compound was not commercially available.

NMR was used for quantitative wax ester analysis as described by Santala et al. (2011). Briefly, samples were prepared by collecting cell pellets from 40 ml of cell culture by centrifugation and freeze-drying the cell pellet (ALPHA 1–4 LD plus freeze dryer, Martin Christ, Germany). Cell dry weight (CDW) was measured form the freeze-dried cell pellets before lipid extraction for ^1^H NMR measurements (Varian Mercury spectrometer, 300 MHz). The samples were dissolved chloroform with trifluorotoluene as an internal standard and the spectra was measured. Data was processed with ACD NMR processor program and interpreted accordingly. This NMR method quantifies α*-*alkoxy methylene protons of ester bonds that are specific to wax esters.

#### Results and Discussion

*A. baylyi* ADP1 belongs to a bacterial species capable of metabolizing a variety of aromatic compounds by enzymatically funneling them towards central intermediates. The compounds are metabolized via the β-ketoadiapate pathway to succinyl-CoA and acetyl-CoA (Wells and Ragauskas, 2012). As the latter is an essential intermediate for the synthesis of fatty-acid derived products, such as the naturally produced wax esters in *A. baylyi* (Reiser and Somerville, 1997), (Stöveken and Steinbüchel, 2008) (Uthoff and Stöveken, 2005), the aromatic-catabolizing pathway could potentially provide means to utilize lignin-derived molecules for the production of long-chain carbon compounds. We investigated the ability of *A. baylyi* ADP1 to utilize lignin-derived molecules for wax ester and alkane synthesis, and determined how structural differences of the selected aromatic compounds affect biomass and product synthesis. We employed two ADP1 strains described previously: The first one, designated here as ‘WP’, synthesizes wax esters *via* the native synthesis pathway (Santala et al., 2011). The strain exhibits a previously described bioluminescence based - sensor for the detection of long-chain aldehydes, that are specific intermediates in the wax ester synthesis pathway (Lehtinen et al., 2018a; Santala et al., 2011). The second strain, designated here as ‘AP’, is engineered for the synthesis of alkanes by a non-native pathway, and contains a sensor system for the detection of both aldehydes (a precursor in alkane synthesis) monitored via bioluminescence, and alkanes, detected as fluorescence (Lehtinen et al., 2017b). To avoid degradation of the produced alkanes or direction of acetyl-CoA to wax esters instead of alkanes, the native alkane-degrading pathway and the wax ester synthesis pathway of *A. baylyi* ADP1 have been disrupted. The proposed carbon flow from substrate to product in the strains WP and AP is presented in Figure 1.

**Figure 1:**
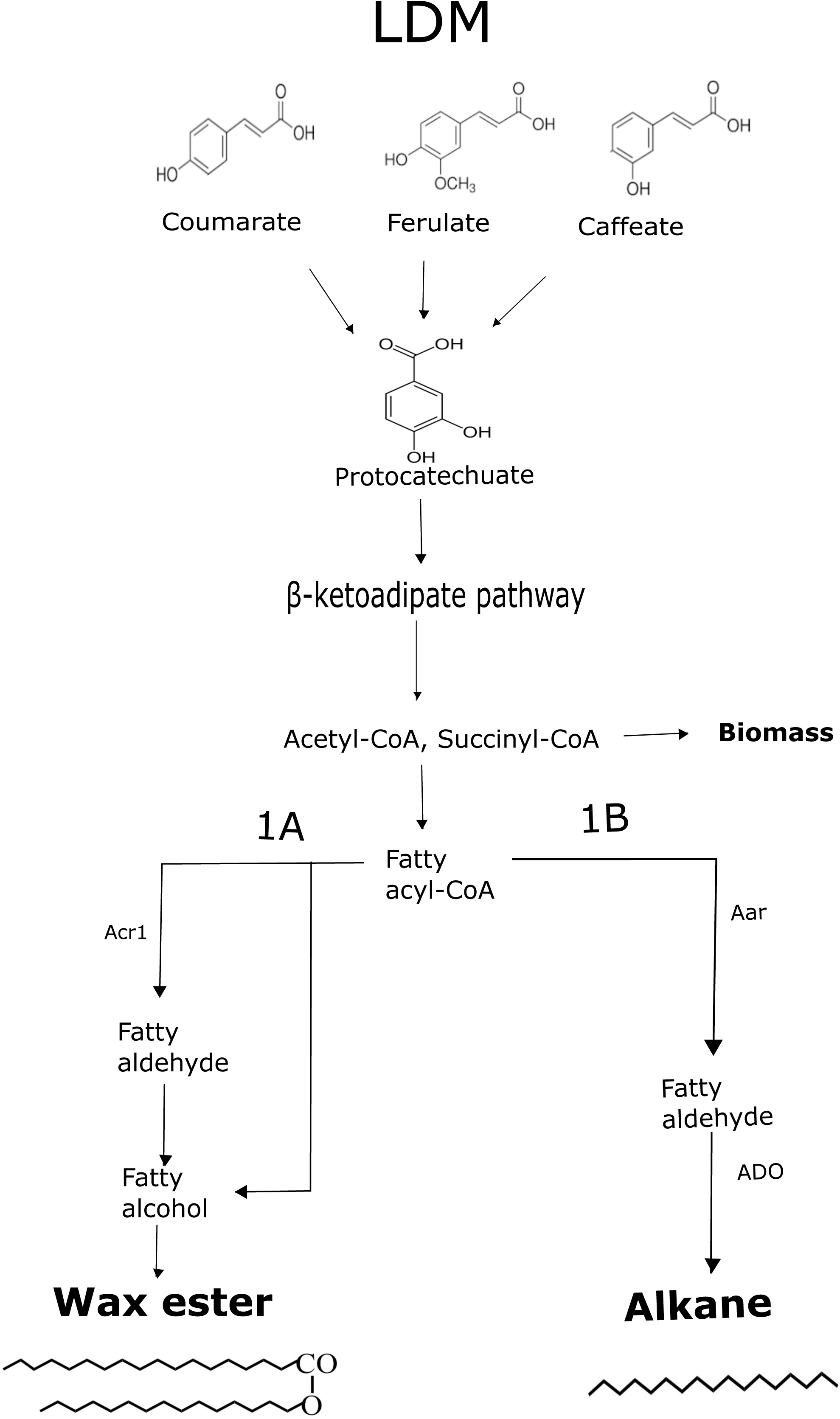
The schematic presentation of carbon flow from lignin-derived monomers (LDMs; coumarate, ferulate, caffeate) into products. The structurally analogous LDMs are first funneled into a single intermediate, protocatechuate. After ring cleavage, the intermediate is metabolized by the β-ketoadipate pathway yielding acetyl-CoA and succinyl-CoA. From acetyl-CoA, two different pathways for possible products (1A for wax esters, 1B for alkanes) are shown. In the native wax ester synthesis pathway, first the fatty acyl-CoA is produced and then fatty alcohols *via* a fatty aldehyde intermediate. The fatty alcohols are esterified with fatty acyl-CoAs resulting in wax esters. In the alkane producing strain, the natural fatty aldehyde reductase gene *acr1* has been replaced with a non-native reductase gene, *aar*. The fatty aldehydes produced by AAR are further converted to alkanes by another heterologous enzyme, aldehyde deformylating oxygenase, ADO.

### Product synthesis and biomass from LDM representatives

The WP and AP strains were employed to study the potential of coumarate, ferulate and caffeate as carbon sources for simultaneous product synthesis and biomass formation. Cultivations were carried out by supplementing 15 mM of coumarate, ferulate or caffeate together with 25 mM of acetate. The WP strain utilized both coumarate and ferulate efficiently, although product formation measured as luminescence signal was higher from coumarate (Figures 2A, 2B and 2C). Biomass, on the other hand, was produced rather similarly between these two compounds. In both cases, growth ceased within 20 hours followed by a rapid drop in the luminescence signal (Figure 2B). Caffeate supplementation, on the other hand, revealed possible inhibition by the compound seen as prolonged lag phase, and low luminescence signal and biomass formation (Figures 2A, 2B and 2C). However, ADP1 wild type has previously been shown to consume caffeate at a lower concentration of 10 mM (Salmela et al., 2018b).

**Figure 2.**
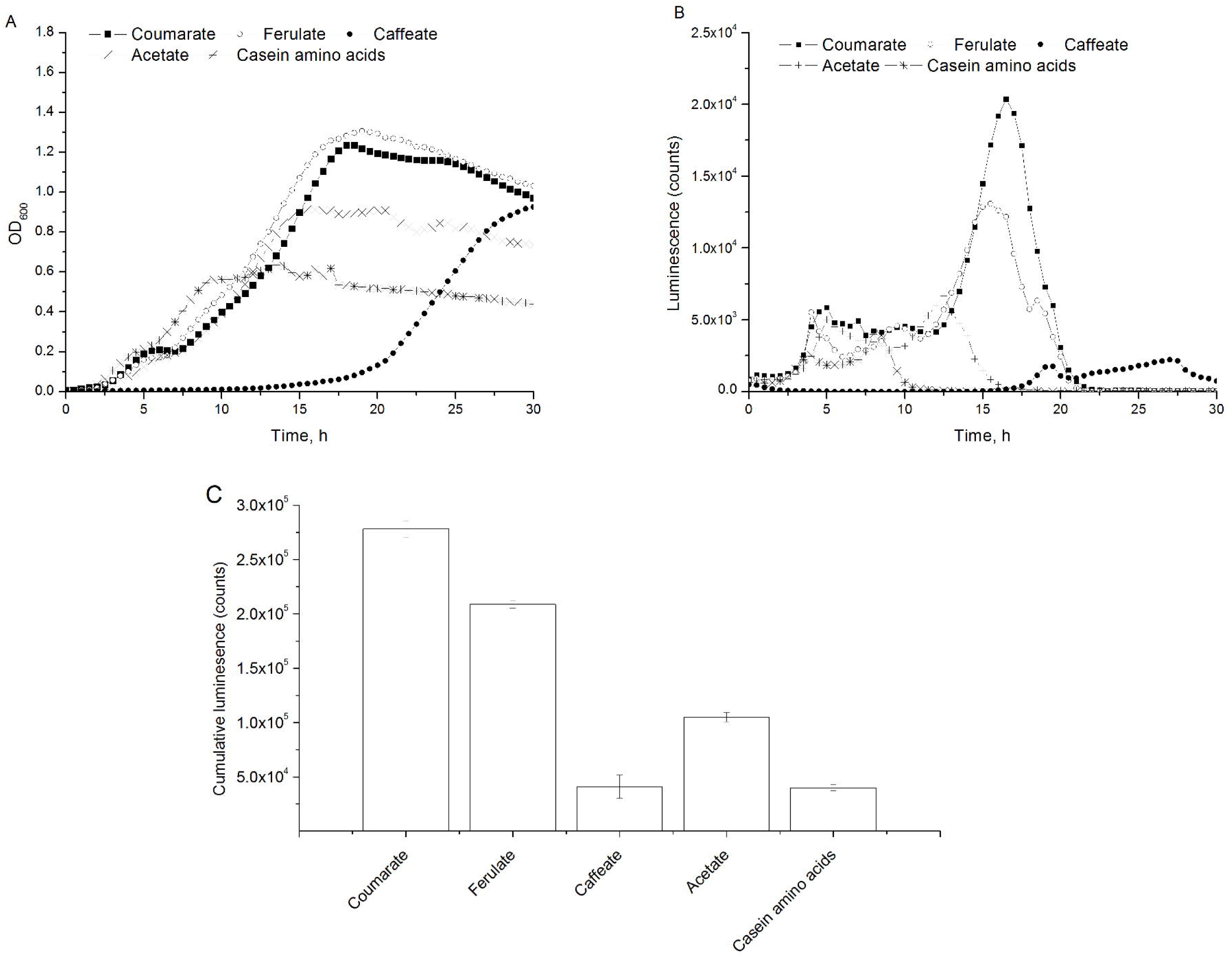
Biomass and product formation by the WP strain utilizing different LDMs as substrates. Error bars have been left out from the A) and B) for clarity and are available in supplementary material (Figure S1). **A)** Cell growth measured as optical density at 600 nm every 30 minutes. Carbon sources used: 25 mM acetate and 0.2% casein amino acids with 15 mM coumarate (closed square), ferulate (open circle) or caffeate (closed circle) supplementation. Control cultivations supplemented with 25 mM acetate and casein amino acids (cross) or casein amino acids (star). The results are the average of three biological replicates. **B)** Real-time luminescence signal representing the internal aldehyde (wax ester precursor) formation measured every 30 minutes. The results are the average of three biological replicates. **C)** Cumulative luminescence signals representing relative product formation from the different LDMs. The results are the average of three biological replicates and error bars represent standard deviation.

The optical density, luminescence and fluorescence profiles of the AP strain show similar substrate preferences as with the WP strain; coumarate and ferulate were efficiently utilized for simultaneous biomass and alkane production (Figures 3A, 3B, 3C and 3D). Although caffeate supplementation yielded somewhat better growth for the AP strain than with the WP strain (Figure 3A), low luminescence and fluorescence signals indicate inefficient direction of substrate for the product formation (Figures 3c and 3D). Estimated by the fluorescence signal, alkane production was highest with coumarate as the carbon source (Figure 3D). This correlates with the higher cumulative luminescence obtained from coumarate-supplemented cultivations compared to the other carbon sources (3C). Furthermore, the cumulative luminescence signal produced by the AP strain from coumarate was 11.5 fold higher than with the WP strain. The heterologous enzyme AAR is more efficient in supplying aldehyde substrate for the bacterial luciferase LuxAB, which explains the higher obtained luminescence signal (Lehtinen et al., 2017b; Lehtinen et al., 2018a). Thus, the architecture and the comprising enzymes of a production pathway may have a significant role in determining the production yield in the context of LDM utilization.

**Figure 3.**
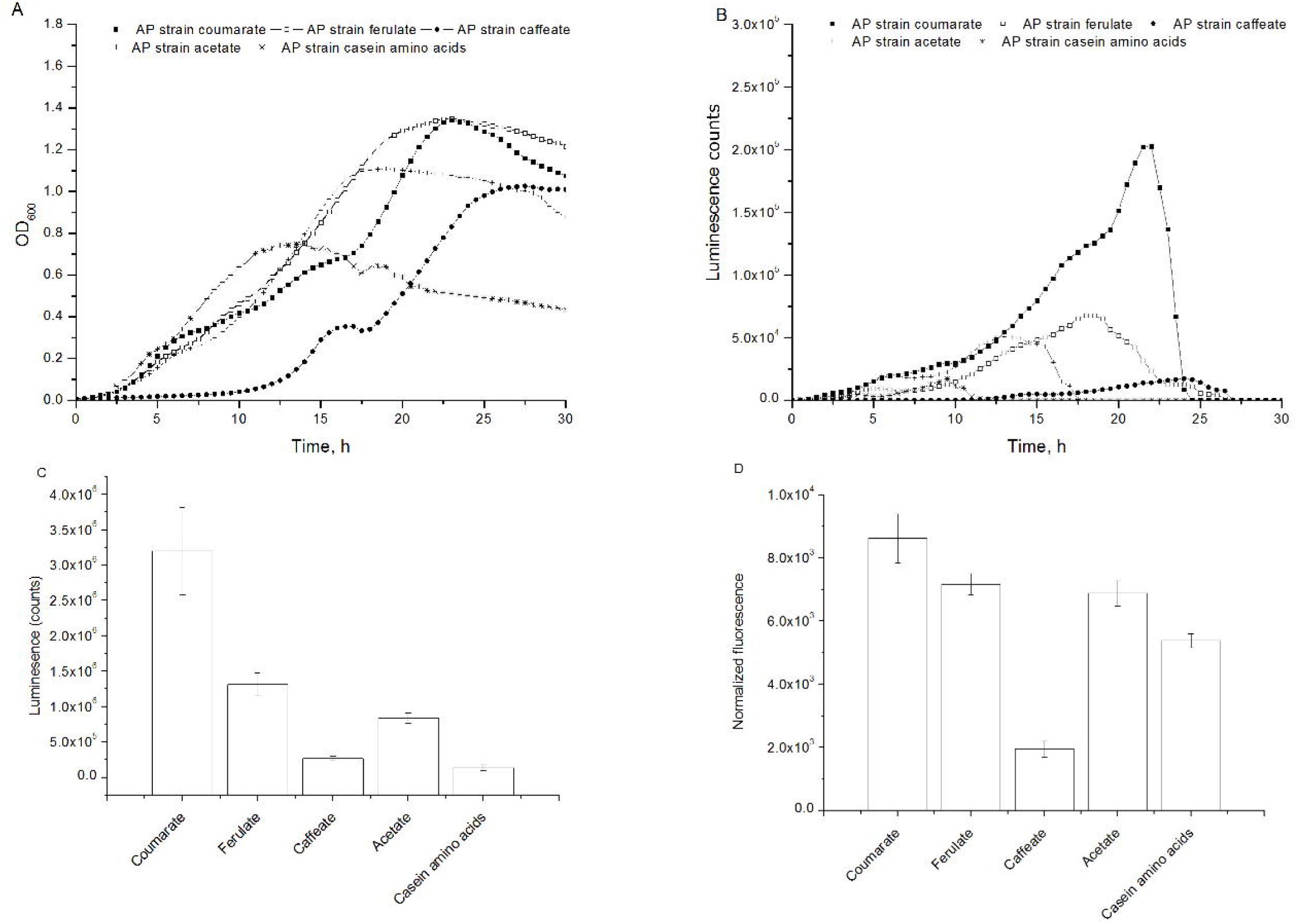
Biomass and product formation by the AP strain utilizing different LDMs as substrates. Error bars have been left out from the A) and B) for clarity and are available in supplementary material (Figure S1). **A)** Cell growth measured as optical density at 600 nm every 30 minutes. Carbon sources used: 25 mM acetate and 0.2% casein amino acids with 15 mM coumarate (closed square), ferulate (open circle) or caffeate (closed circle) supplementation. Control cultivations supplemented with 25 mM acetate and casein amino acids (cross) or casein amino acids (star). The results are the average of three biological replicates. B) Real-time luminescence signal representing the internal aldehyde (alkane precursor) formation measured every 30 minutes. The results are the average of three biological replicates. C) Cumulative luminescence signals representing relative product formation from the different LDMs. The results are the average of three biological replicates and error bars represent standard deviation. **C)** Cumulative luminescence signals representing relative product formation from the different LDMs. The results are the average of three biological replicates and error bars represent standard deviation. **D)** Normalized fluorescence signals (fluorescence/OD_600_) representing alkane production from LDMs. The results are the average of three biological replicates and error bars represent the standard deviation of the samples.

The biochemistry of the funnelling pathway in *A. baylyi* is similar to those of the other β-ketoadipate utilizing microorganism, such as *Pseudomonas putida* (Parke et al., 2000)(Harwood and Parales, 1996). In both microorganisms, LDMs such as coumarate, ferulate and caffeate are converted through the protocatechuate branch to single intermediates. The biocatalytic efficiencies of the first enzymatic steps of this upper funneling pathway may vary between the carbon sources. For example, muconate production by a genetically engineered *P*. putida KT2440-CJ102 yielded slightly higher conversion rates for coumarate than ferulate (Johnson et al., 2017). The structural differences between the studied LDMs might explain why coumarate is favored over ferulate and caffeate. Ferulate requires demethylation of the aromatic structure, whereas the aromatic ring structure of coumarate can be directly hydroxylated to the central intermediate protocatechuate. 4-HBA and vanillate are produced as overflow metabolites from coumarate and ferulate (Parke and Ornston, 2003), and although not yet fully elucidated in *A. baylyi*, the transport efficiencies of these compounds may also vary. Although real lignin-derived streams are more complex in their composition than these model compounds, studies on individual compounds reveal valuable details about the host metabolism for future process design.

### Tolerance to combined effects of coumarate and acetate

Microorganisms with high tolerance towards inhibitory compounds, such as aromatics and acetate, are attractive candidates for the possible upgrading of lignin-derived molecules to value added products. Acetate has been associated as a part of hardwood lignins (Lu and Ralph, 2010), as well as a residual components produced by lignocellulose treatment processes (Jönsson and Martín, 2016). Previously, *A. baylyi* has been shown to tolerate and utilize acetate as a sole carbon source at concentrations as high as 200 mM (Lehtinen et al., 2017b). Complex substrate regulation systems in *A. baylyi* include global regulators causing, for example, carbon-catabolite repression, as well as vertical and horizontal regulation by intermediates (Bleichrodt et al., 2010). For example, *A. baylyi* ADP1 prefers acetate as a carbon sources over the aromatic compounds (Zimmermann et al., 2009). Consequently, this regulation may result in sequential use of the carbon sources, slower conversion rates, and eventually in inefficient product formation and substrate utilization. On the other hand, reactions occurring at the β-ketoadipate pathway are oxygenase mediated, and do not provide ATP reserves (Ornston and Stanier, 1966). Thus, an alternative carbon source typical for lignin processes, such as acetate, provides means to accelerate cellular growth.

To determine the effect of acetate on coumarate utilization and to elaborate the inhibitory effect of using a mixture of coumarate and acetate, the WP strain was cultured at different concentrations of acetate (25–90 mM) and coumarate (15–50 mM). Coumarate was chosen as a model compound based on the results obtained from the product synthesis experiments with different LDM-representatives. With acetate and coumarate concentrations up to 55 mM and 30 mM, respectively, efficient formation of biomass and products was observed, whereas with 75 mM acetate and 45 mM coumarate, cell viability was severely impaired (Figures 4, 4B). Furthermore, higher substrate concentrations promoted more diauxic growth. In addition to substrate inhibition, elevated concentrations of the metabolic key intermediates, such as carboxymuconate and its precursor protocatechuate, are toxic to cells (Parke et al., 2000). Towards the end of the experiment, subtle growth was also observed with the cultures supplemented with 90 mM acetate and 50 mM coumarate. Although *A. baylyi* tolerates rather high concentrations of acetate and coumarate, lower concentrations allow for faster and more efficient growth and product formation. For comparison, glucose grown *P. putida* KT2240 and *E. coli* MG1655 were inhibited (33% reduction in growth rate) by 61 mM and 30.4 mM of coumarate or 65.6 mM and 91.4 mM of acetate, respectively, when supplemented individually Calero et al. (2017). To obtain higher tolerance towards inhibitory substrates, strain optimization could be applied through adaptation and laboratory evolution (Dragosits and Mattanovich, 2013) or by genetic engineering. For example, Benndorf et al. (2001) showed that by inducing *A. calcoaceticus* 69-V (*A. baylyi* previously referred to as *A. calcoaceticus*) with 14 mM of phenols or catechols heat shock proteins were produced to increase tolerance towards oxidative stress. Kohlstedt et al. 2018 achieved a 20% higher tolerance towards catechols in *P. putida* by expressing native catA and catA2 genes under the same catA promoter.

**Figure 4:**
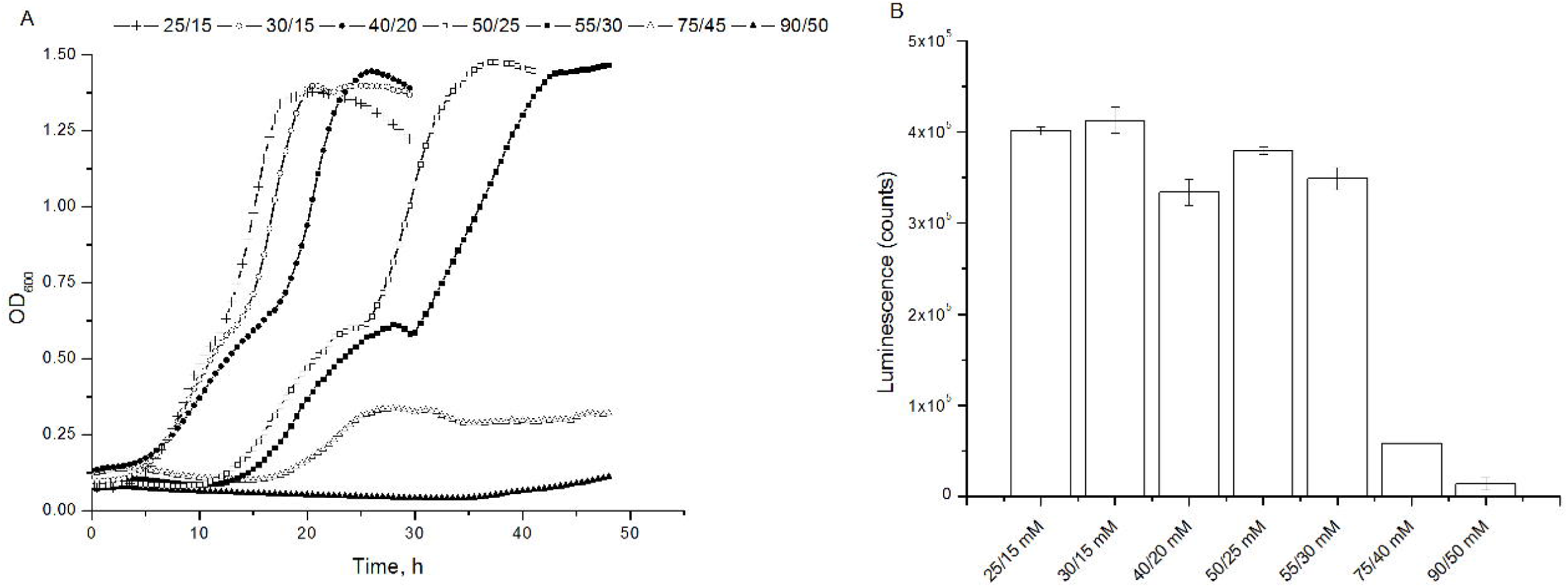
The effect of increasing acetate and coumarate concentrations on growth and aldehyde formation by the WP strain cultivated for 48 hours. **A)** Growth profiles as optical density (OD_600_) measured every 30 minutes. Substrate concentrations used: 25mM acetate and 15 mM coumarate (cross), 30 mM acetate and 15 mM coumarate (open circle), 50mM acetate and 25 mM coumarate (opens square) 55 mM acetate and 30 mm coumarate (closed square), 75 mM acetate and 40 mM coumarate (open triangle) and 90 mM acetate and 50 mM coumarate (closed triangle). **B)** Cumulative luminescence signal from the different carbon sources at the end of the experiment (48 h). Error bars represent the standard deviation of three biological replicates. Error bars have been left out for clarity and are available in supplementary material (Figure S2).

#### Wax ester and alkane production from coumarate in batch cultures

Based on the previous experiments, coumarate was used as the carbon source for product level investigation of wax esters and alkane accumulation from LDMs. Products were analyzed after 8, 12 and 24 h from 50 ml cultivations supplemented with 25 mM of acetate and 15 mM of coumarate. Casein amino acids or 25 mM of acetate and casein amino acids supplementation were used as control cultivations. As hypothesized, the WP strain was able to produce wax esters from coumarate (Figure 5, Table 1). Both acetate and coumarate were consumed rather simultaneously (Table 1) in the studied conditions, although 6.4±0.6 mM 4-HBA had accumulated at the time point of 12 h as an intermediate from coumarate conversion. Experimental data on substrate preferences of *A. baylyi* wild type (Supplementary Figure S3) show that 4-HBA–the overflow metabolite from coumarate— is depleted from the media mainly after all other carbon sources have been depleted. Similarly, after 24 hours, 4-HBA was depleted by the WP strain resulting in a significantly higher biomass compared to the control cultivations. Cell death caused by carbon depletion was observed as decreased OD_600_ after 24 hours. The wax ester content increased during the first 12 h, whereas after 24 h, a weaker wax ester band on TLC was observed (Figure 5). In the acetate control cultivations, the wax esters were depleted already within 12 h, whereas in the casein amino acid control cultivations wax esters were not detected. The depletion of wax esters in carbon deficient conditions has been recorded for example by Fixter et al. (1986) and Santala et al. (2011). Therefore, using substrates such as LDM pose a challenge for wax ester production and preservation due to their inhibitory effects at elevated concentrations. Unspecific strategies, such as adaptive laboratory evolution could allow the use of higher substrate concentrations, whereas metabolic engineering can provide means to circumvent the challenges related storage compound degradation (Lehtinen et al., 2017b).

**Table 1:** Carbon source depletion presented as consumed substrate per supplemented substrate (%) and growth as OD_600_ by the WP strain measured at three time points (8, 12 and 24h). Batches were supplement either with acetate, coumarate and casein amino acids, acetate and casein amino acids or casein amino acids. Results are presented for three biological replicates with standard deviation marked as ±.

|  | 25 mM acetate, 15 mM coumarate and 0.2% casein amino acids |  |  | 25 mM acetate and 0.2% casein amino acids |  | 0.2% casein amino acids |
| --- | --- | --- | --- | --- | --- | --- |
| Time, h | Coumarate consumed, % | Acetate consumed, % | OD | Acetate consumed, % | OD | OD |
| 8 | 28.7±3.5 | 45.0±3.5 | 1.60±0.10 | 88±14 | 1.49±0.27 | 0.16±0.03 |
| 12 | 100.0 | 100.0 | 3.43±1.81 | 100 | 1.45±0.08 | 0.19±0.02 |
| 24 | 100.0 | 100.0 | 2.01±0.17 | 100 | 0.66±0.34 | 0.16±0.01 |

**Figure 5.**
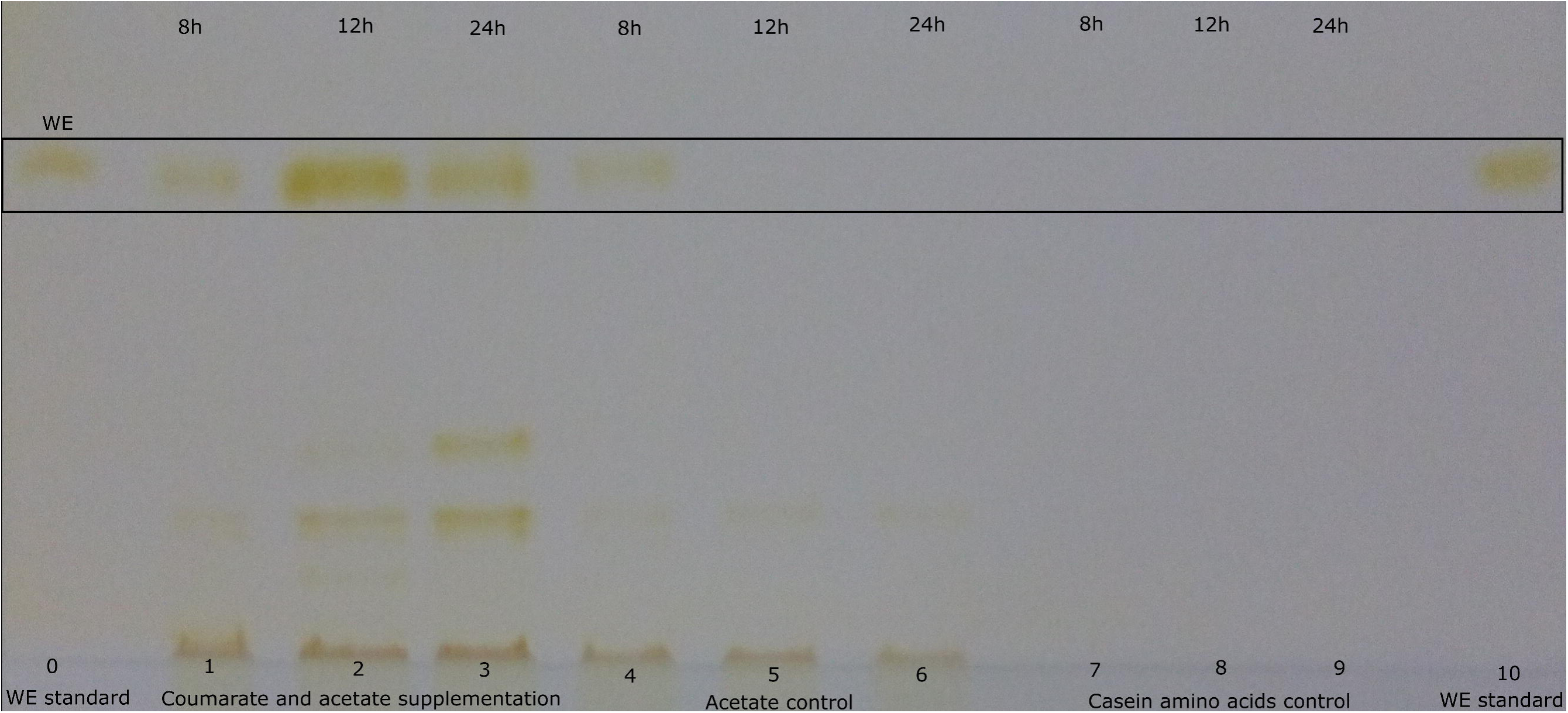
Semi-quantitative TLC analysis of the wax esters produced by the WP strain at different time points. Samples were grown on 25 mM acetate, 15 mM coumarate and 0.2% casein amino acids (lanes 1, 2 and 3), 25 mM acetate and 0.2% casein amino acids (lanes 4, 5 and 6) or 0.2% casein amino acids (lanes 7, 8 and 9). Lanes 0 and 10 represent the wax ester standard (Jojoba oil). Time points for each sample (8h, 12h and 24h) are shown at the top of the figure.

In the AP strain, the gene responsible for alkane degradation has been knocked-out, resulting in an intracellular product that is non-degradable by the host metabolism. Here, we demonstrated that the AP strain produced alkanes from coumarate in a 12-hour batch cultivation. In the culture, acetate and coumarate were depleted within 8 hours, although 11 mM of 4-HBA had accumulated during this time (Table 2). Already at this time point, heptadecane was detected (Figure 6). After 12 hours, 4-H BA was consumed and a significant increase to 169 μg/L of heptadecane was observed, whereas production from acetate and casein amino acids control and casein amino acid control was negligible, 16.8μg/L and 12.0 μg/L respectively. After 24 hours, an intriguing phenomenon was observed when analyzing the alkanes: although the heptadecane concentration slightly decreased towards the end of the experiments, the amount of 8-heptadecene increased indicating that double bond conversion may have taken place inside the cells (Supplementary Figure S3). Furthermore, it is possible that some of the alkanes were lost in the supernatant due to carbon starvation and cell lysis. Heptadecane was produced modestly in the batch experiments: the C/C yield from acetate and coumarate was slightly higher (0.006%) when compared to our previous studies (0.005%) using acetate as a sole substrate (Lehtinen et al., 2018b). Furthermore, the heptadecane yield from consumed coumarate was estimated to be 62.3 μg/g. It should be noted that coumarate serves as a substrate for both biomass formation and product synthesis. For efficient direction of carbon to product, decoupling of product and biomass synthesis could be achieved by metabolic engineering (Santala et al., 2018).

**Figure 6.**
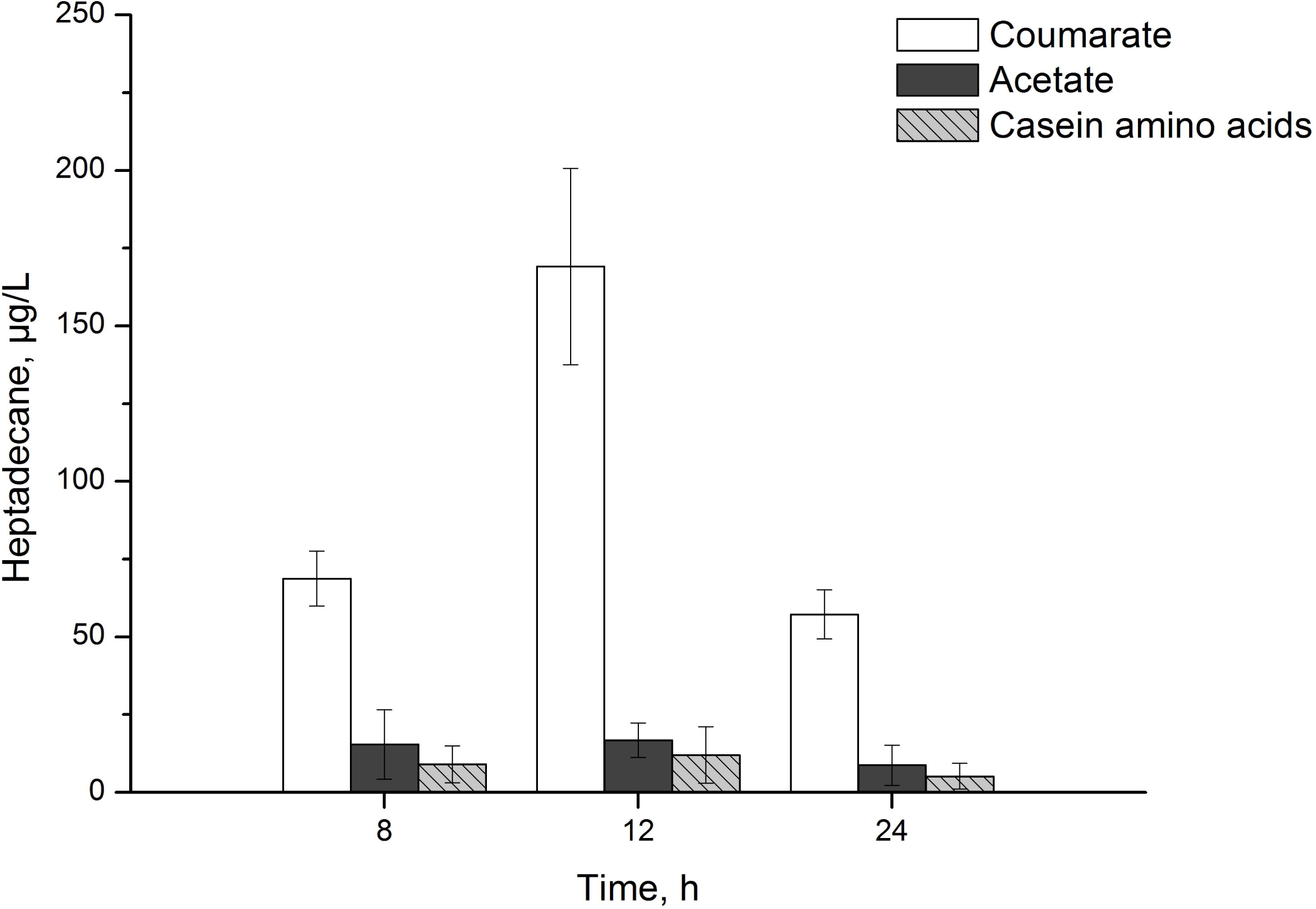
Heptadecane production as μg/L by the AP strain from 25 mM coumarate, 15 mM acetate and 0.2% casein amino acids (white columns), 25 mM acetate and 0.2% casein amino acids (grey columns) and 0.2% casein amino acids (striped grey columns) at 8h, 12 h and 24h. The error bars represent the standard deviation from three biological replicates.

**Table 2:** Carbon source depletion presented as consumed substrate per supplemented substrate (%) and growth as OD_600_ by the AP strain measured at three time points (8, 12 and 24h Batches were supplement either with acetate, coumarate and casein amino acids, acetate and casein amino acids or casein amino acids. Results are presented for three biological replicates with standard deviation marked as ±.

|  | 25 mM acetate, 15 mM coumarate and 0.2% casein amino acids |  |  | 25 mM acetate and 0.2% casein amino acids |  | 0.2% casein amino acids |
| --- | --- | --- | --- | --- | --- | --- |
| Time, h | Coumarate consumed, % | Acetate consumed, % | OD | Acetate consumed, % | OD | OD |
| 8 | 94.7±0.5 | 82.8±4.35 | 1.87±0.13 | 100 | 1.43±0.0.22 | 0.12±0.03 |
| 12 | 100.0 | 100.0 | 2.51±0.17 | 100 | 1.12±0.08 | 0.11±0.02 |
| 24 | 100.0 | 100.0 | 2.10±0.06 | 100 | 0.66±0.03 | 0.16±0.01 |

### Wax ester production from coumarate in bioreactor

The WP strain produced wax esters from coumarate effectively in the 50 ml batch studies. Thus, a bioreactor experiment was conducted to elucidate the dynamics of wax ester accumulation from coumarate. One of the challenges in maintaining reactions favorable towards product formation when utilizing LDMs, such as coumarate, is caused by substrate inhibition. With low substrate doses, swift carbon depletion is followed by rapid wax ester degradation. Interestingly, in our shake-flask cultivations wax esters were detected even after prolonged carbon starvation. As the ring cleavage of aromatic compounds by bacteria is an oxygen intensive process (Fuchs et al., 2011), we tested whether the premature wax ester degradation could be avoided by restricting the metabolic activity of the cell by limiting the oxygen supply in the bioreactor. Furthermore, we fed the coumarate gradually in the bioreactor to study the effect of substrate concentration for the wax ester production.

An initial total volume of 750 ml medium was supplemented with 25 mM of acetate and 15 mM of coumarate and inoculated with the WP strain. After substrate depletion, the reactor was re-supplemented twice with elevated coumarate concentrations (25 mM and 34 mM). Automated supply of pure oxygen (1 vvm) was initiated when the pO_2_ decreased below 20%. Regardless of the additional O_2_ supply, the pO_2_ remained at 0% throughout coumarate utlization and spiked only at substrate depletion (Figure 7). Both acetate and coumarate were consumed during the first 10 hours of cultivation and wax esters accumulated at time points of 7, 8 and 9 hours (Figure 7) up to 26 mg/L. However, after carbon depletion, the wax esters were consumed rapidly within 60 minutes at time point of 10 hours.

**Figure 7:**
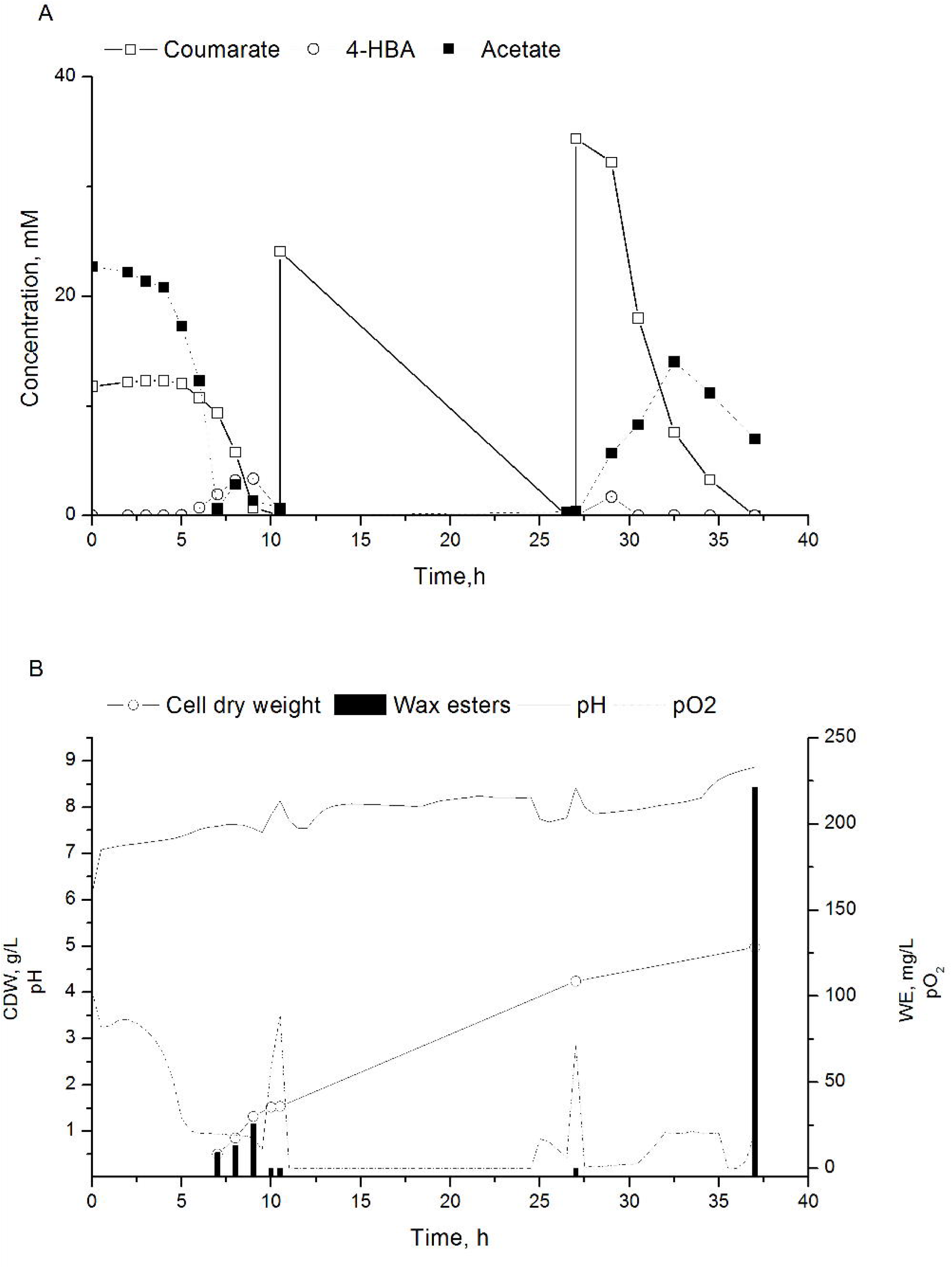
Bioreactor experiment for wax ester accumulation by the WP strain. **A)** Substrate concentrations (mM) of coumarate (open square), 4-HBA (open circle) and acetate (closed square) during 37 hours of cultivation. Initially, the reactor was supplemented only with acetate and coumarate. At time points 10.5 h and 26.5 h, the cultivations were re-supplemented with coumarate. 4-HBA is produced as an intermediate of coumarate conversion. **B)** Wax ester titer as mg/L (columns), biomass formation as cell dry weight in g/L (open circles), partial oxygen pressure as percentage (dotted line) and pH (line) during 37 hours of cultivation. Please note, that the right y-axis starts at −5 to indicate wax ester content of 0 mg/l at time points 10, 10.5 and 26.5 h.

To study the effect of oxygen in wax ester degradation, the pure oxygen supply was shut off at 12 h and the reactor was aerated only with an airflow at a rate of 1 vvm. At this point, 25 mM of coumarate was also added to the reactor. After 12 h cultivation in these conditions, the supplemented coumarate had been depleted and cell growth observed, though wax esters were not detected (Figure 7). The automated O_2_ supply was re-initiated and an elevated concentration of coumarate was added to the reactor. Re-supplementation with an increased coumarate concentration (34 mM) resulted in rapid utilization of coumarate for biomass and wax esters between the time points of 27–37 h. Small amounts of 4-HBA accumulated during these 10 hours. In addition, a peak in acetate accumulation (14 mM) was observed at time point of 29 hours. At the end of the experiment at 37 hours, the obtained biomass was 5.0 g/L. The pH of the cultivation varied between pH 6.1–8.3 until finally reaching 8.7 at the end of cultivation. Intracellular wax esters were produced up to 221 mg/L at the end of the bioreactor experiment and a yield of 40 mg_wax ester_/g_coumarate_ was obtained between time points 27–37. The yield is higher than previously obtained results for *A. baylyi* wild type (14–20 mg_wax ester_/g_substrate_) when using glucose or acetate as a substrate (Lehtinen et al., 2018b)(Santala et al., 2018) and implicates coumarate as a potential carbon source for wax ester production.

## Conclusions

The ability of microorganisms to produce industrially relevant compounds from low cost substrates, as well as the flexibility of modifying the cellular systems to produce non-native products, pave the way for a more sustainable biobased economy. Here, we showed that LDMs can be used for long chain alkyl ester (C_32_-C_34_) production – the naturally accumulated storage compounds of *A. baylyi –* as well as for the production of drop-in fuel components in the form of long chain alkanes (C_17_) produced by a synthetic pathway. In addition, we observed that the chemical structure of the studied LDMs affect biomass and product synthesis, coumarate being the most propitious for product and biomass formation. Thus, the choice of biomass and pre-treatment methods could be adjusted to generate LDMs that are optimal for production.

---

**Supplementary Figure S1**. Optical densities and luminescence counts from Figures 2A, 2B, 3A and 3B presented with error bars. Error bars represent the standard deviation from three biological replicates.

**Supplementary Figure S2**. Optical densities from Figure 4 presented with error bars. Error bars represent the standard deviation from three biological replicates.

**Supplementary Figure S3**. *A. baylyi* wild type substrate consumption in a 50 ml batch cultivation supplemented with 25 mM acetate (black square) and 15 mM coumarate (open square). 4-HBA (black diamond) accumulates as an overflow metabolite from coumarate conversion. Cell growth measured as OD_600_ (open circle).

**Supplementary Figure S3**. GC-MS chromatogram showing the increase of 8-heptadecene (RT 11.8) and decrease in heptadecane (RT 11.9). Blue line t=24h, black line t=12.

