## Supplemental Figures for "Alkane and wax ester production from lignin derived molecules"

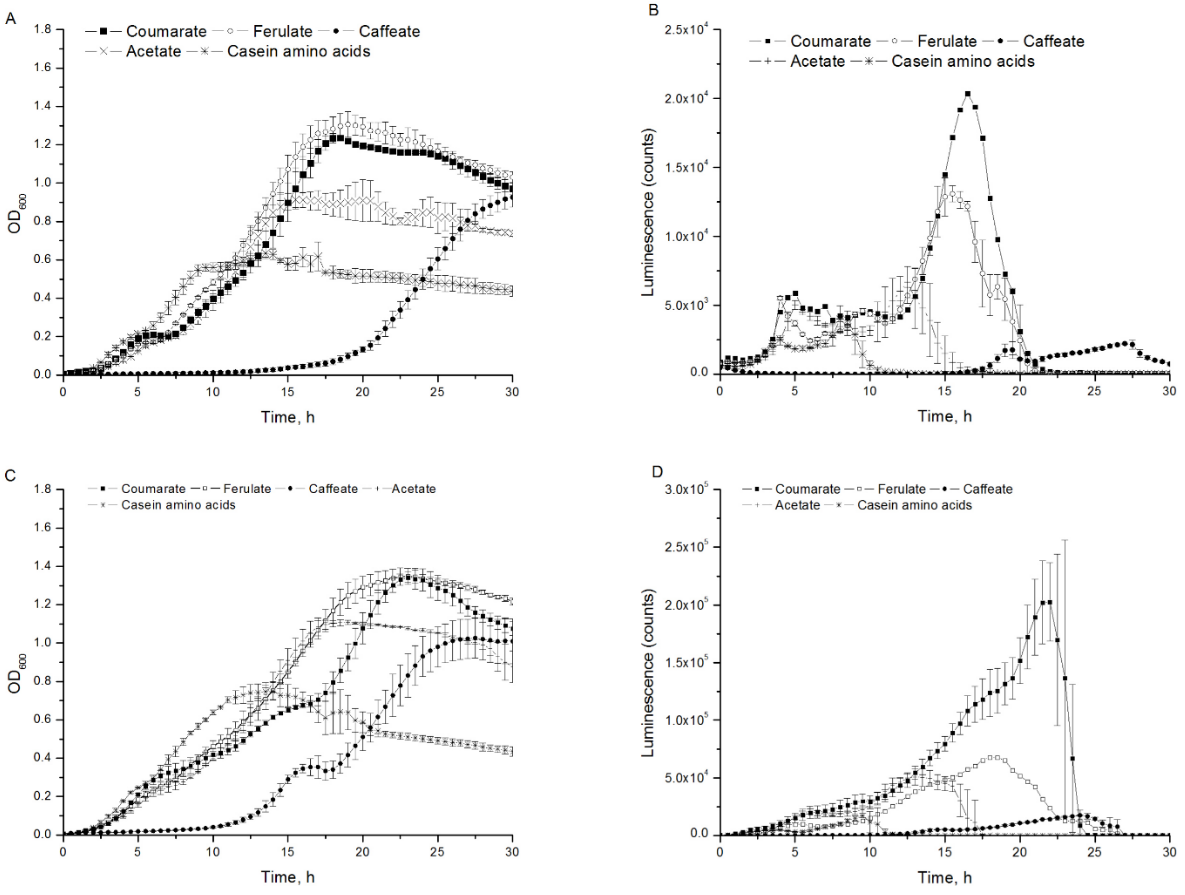


**Supplementary Figure S1.** Optical densities and luminescence counts from Figures 2A, 2B, 3A and 3B presented with error bars. Error bars represent the standard deviation from three biological replicates.

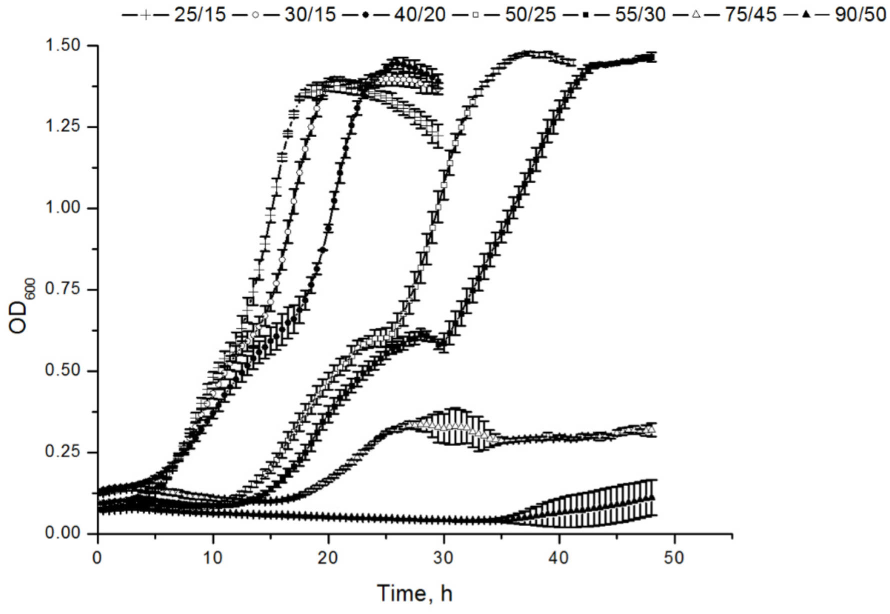


**Supplementary Figure S2.** Optical densities from Figure 4 presented with error bars. Error bars represent the standard deviation from three biological replicates.

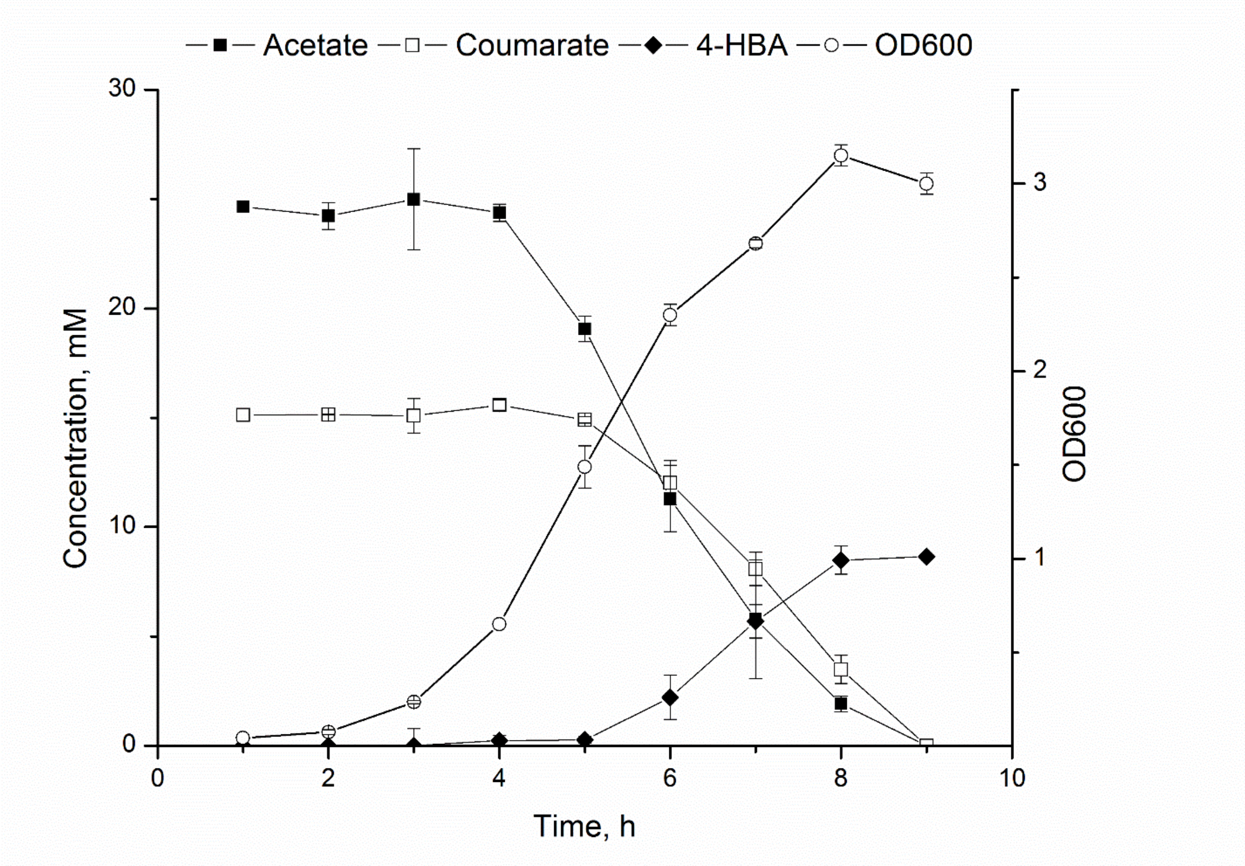


**Supplementary Figure S3.** *A. baylyi* wild type substrate consumption in a 50 ml batch cultivation supplemented with 25 mM acetate (black square) and 15 mM coumarate (open square). 4-HBA (black diamond) accumulates as an overflow metabolite from coumarate conversion. Cell growth measured as $OD_{600}$ (open circle).

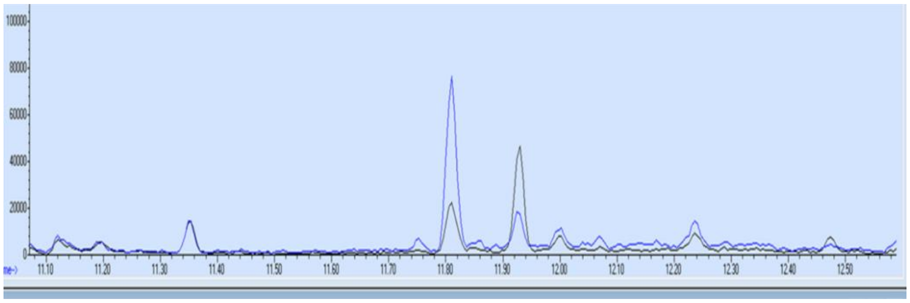

**Supplementary Figure S3.** GC-MS chromatogram showing the increase of 8-heptadecene (RT 11.8) and decrease in heptadecane (RT 11.9). Blue line t=24h, black line t=12.
